# Imaging-based screening identifies modulators of the *eIF3* translation initiation factor complex in *Candida albicans*

**DOI:** 10.1101/2023.04.19.537517

**Authors:** Katura Metzner, Matthew J O’Meara, Benjamin Halligan, Jesse W. Wotring, Jonathan Z Sexton, Teresa R O’Meara

**Affiliations:** Department of Microbiology and Immunology, University of Michigan Medical School, Ann Arbor, MI, USA; Department of Computational Medicine and Bioinformatics, University of Michigan, Ann Arbor, MI, USA; Department of Medicinal Chemistry, College of Pharmacy, Ann Arbor, MI, USA; University of Michigan Center for Drug Repurposing, USA; Department of Internal Medicine, Gastroenterology, University of Michigan Medical School, Ann Arbor, MI, USA

## Abstract

Fungal pathogens like *Candida albicans* can cause devastating human disease. Treatment of candidemia is complicated by the high rate of resistance to common antifungal therapies. Additionally, there is host toxicity associated with many antifungal compounds due to the conservation between essential mammalian and fungal proteins. An attractive new approach for antimicrobial development is to target virulence factors: non-essential processes that are required for the organism to cause disease in human hosts. This approach expands the potential target space while reducing the selective pressure towards resistance, as these targets are not essential for viability. In *C. albicans,* a key virulence factor is the ability to transition to hyphal morphology. We developed a high-throughput image analysis pipeline to distinguish between yeast and filamentous growth in *C. albicans* at the single cell level. Based on this phenotypic assay, we screened the FDA drug repurposing library of 2,017 compounds for their ability to inhibit filamentation and identified 33 compounds that block the hyphal transition in *C. albicans* with IC_50_ values ranging from 0.2 to 150 µM. Multiple compounds showed a phenyl vinyl sulfone chemotype, prompting further analysis. Of these phenyl vinyl sulfones, NSC 697923 displayed the most efficacy, and by selecting for resistant mutants, we identified *eIF3* as the target of NSC 697923 in *C. albicans*.

## Introduction

Fungi have a devastating impact on human health, and the treatment of invasive fungal infections is notoriously difficult. As the number of immunocompromised and hospitalized patients vulnerable to fungal infections increases (1), it is essential to discover new targets and approaches for treating these deadly fungal pathogens. The discovery of antifungals with selective toxicity towards fungi has been hampered by the close evolutionary relationship between fungi and humans (2, 3). In the clinic, there are currently only three classes of antifungals used for the treatment of invasive fungal infections, compared to over two dozen classes for bacterial infections (4, 5). These include the azoles, which target lanosterol 14α-demethylase encoded by the essential gene *ERG11;* amphotericin, which targets ergosterol in the fungal cell membrane; and the echinocandins, which target β-(1,3) glucan synthase encoded by the essential gene *FKS1* (5). However, resistance to each class of antifungal has emerged, thus demanding new therapeutic strategies (6).

To address this issue, a promising strategy is to identify key virulence factors and target those for new antimicrobial therapies (7, 8). Targeting virulence factors also opens new opportunities for chemical diversity in therapeutics, as these molecules are not typically explored by conventional antifungal strategies. One hypothesis is that the genes governing fungal virulence are more likely to be species specific and less conserved between fungi and their mammalian hosts. Transitions between morphological states in response to entry into the human host is a broadly conserved trait amongst fungal pathogens (9). For *Candida albicans*, a major virulence factor is the ability of this organism to transition from yeast to filamentous growth (10–14). Filamentation is linked with tissue invasion, secretion of the Candidalysin toxin, biofilm formation, and induction of macrophage pyroptosis, all of which are important for this organism to cause disease (12, 13, 15–17).

Due to the importance of filamentation during *C. albicans* pathogenesis, there has been intense research on the genetic circuitry required for *C. albicans* to transition between morphological states (12, 13). Despite this, there has not been a corresponding comprehensive analysis of small molecule inhibitors or regulators of this process as the detailed microscopic analysis needed to discriminate between yeasts and filaments is laborious and time-consuming. From previous screens, a few candidate compounds have been identified (8, 18–20), demonstrating the potential of small molecules to regulate *C. albicans* filamentation. However, none of the previous approaches have leveraged the power of high-throughput automated single-cell resolution imaging, thus limiting their capacity to rapidly identify molecules that can inhibit the morphological transition.

Here, we performed a high-content imaging screen that can easily distinguish between yeasts and filaments while simultaneously measuring cell viability. We examined a library of FDA-approved compounds and clinical candidates for their ability to inhibit filamentation, and identified multiple classes of compounds, including steroids, PI3K and TOR inhibitors, and known antifungals. We additionally identified NSC 697923, and two structurally-related compounds, BAY 11-7085 and BAY 11-7082, as inhibitors of filamentation. Bay 11-7082 had been shown to inhibit *C. albicans* biofilms (21, 22), and we hypothesize that this may be through inhibition of filamentation, which is a necessary step in biofilm formation. In mammalian cells, NSC 697923 blocks the formation of a thioester bond between ubiquitin and the active site cysteine of the Ubc13-Uev1A E2 ligase, and BAY 11-7082 and BAY 11-7085 target the Ikk pathway; however, these target proteins are not conserved in *C. albicans.* Therefore, we leveraged a filamentation selection strategy to identify resistant mutants and characterize the mechanism of action of these compounds. Through whole-genome sequencing of resistant mutants, we determined that NSC 697923 compounds may inhibit filamentation through the eIF3 translation initiation complex. Our study suggests that the NSC 697923 chemotype may be repurposed for anti-virulence activity against *C. albicans*.

## Results

### Assay Development

The overarching goal of this project was to discover selective chemical probes that inhibit the yeast to filament transition in *C. albicans*. To develop a robust phenotypic screening platform, we first established a high-content imaging screen that can distinguish between yeasts and filaments in 384-well format with high sensitivity and reliability. We incubated wild type (SC5314) *Candida albicans* cells in YPD with or without 10% serum for four hours at 37 °C as a filament inducing cue (13). Then, both the induced and uninduced cells were fixed, stained with calcofluor white (CFW) and the plates were imaged.

We then developed a CellProfiler image analysis pipeline to accurately segment both yeast and filament cell types (Figure 1A). Briefly, the CFW images were filtered to suppress the appearance of filaments and improve the segmentation of yeast bodies. Yeast were identified as the primary object using Otsu global thresholding and shape-based declumping. Filaments were then segmented as a secondary object using the yeast as the seed object via propagation in the raw CFW image. The resulting yeast and filament regions of interest (ROI) were measured for their size/shape metrics, CFW intensity, and morphologic skeleton; this yielded approximately 300 measurements per cell and an average of 340,000 cells per 384-well plate. To pre-process the cellular features, we first removed zero-variance and location features. Then, to correct for potential differences in staining intensity, we standardized each feature on each plate using the sckit-learn StandardScalar module. We then trained an XGBoost model (v1.6.1) with default parameters over the PC (yeast) and NC (filament) cells. As a quality control, we computed per-plate Z-prime scores. To identify important features driving discrimination, we used SHapley Additive exPlanations (SHAP), as they can handle features that may have non-trivial correlation. We then used the SHAP analysis implemented in the XGBoost library to generate SHAP diagrams and identify high-performer features. To visualize and cluster cells based on their phenotype, we used UMAP to non-linearly embed cell-feature vectors into 2D dimensions. This allowed us to robustly classify cells as either yeast or filaments.

**Figure 1:**
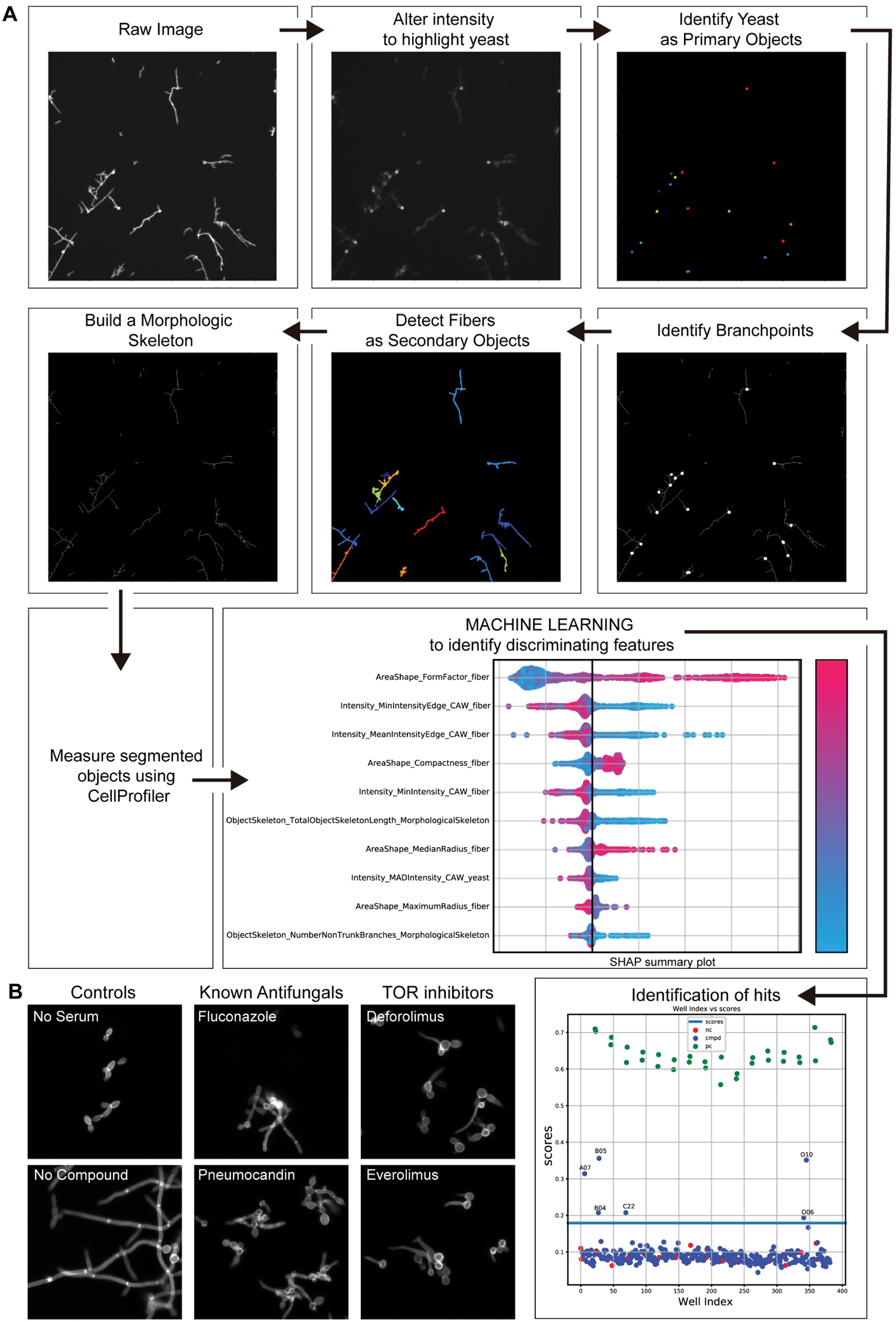
Novel image-based screening approach shows demonstrated ability to identify inhibitors of *Candida albicans* hyphal formation. A) High-throughput microscopy screening assay schematic. B) Representative images of *C. albicans* from the image-based screen. Cells were stained with 1% calcofluor white, and images were taken at 20X magnification.

### Screening

After establishing that the filamentation assay was able to distinguish filaments from yeast, we screened fifteen 384-well plates containing a total of 2,017 unique compounds at a concentration of 10 μM in single wells (Supplemental Table 1). In this screen, we identified 33 compounds that altered *C. albicans* morphology. We were able to capture many known antifungals, including multiple azole and echinocandin drugs, validating our screening approach. If the compounds acted by inhibiting growth, we were also able to use our pipeline to identify compounds that decreased the total number of cells in the well—this was the case for many of the known antifungals, including caspofungin and the antiseptic Gentian Violet. Notably, we identified multiple steroid compounds with an effect on filamentation, including estrone, epiandrosterone, and dehydroepiandosterone (Supplemental Figure 1). Additionally, we identified TOR inhibitors, including everolimus and deforolimus (Figure 1B), consistent with a role for TOR in regulating *C. albicans* filamentation (23). The screening approach also allowed us to identify multiple compounds that inhibited *C. albicans* filamentation without a known mechanism of action.

### Assay Validation

We then performed 10-point dose response assays on the 33 initial hits, and a set of an additional 15 compounds based on chemical similarity or proposed mechanism of action (**Supplemental Table 2**). This analysis of the structurally related compounds was included as it would allow for preliminary structure-activity relationship information. Using this approach, we identified 11 compounds with dose-responsive activity (**Figure 2A**). This set included the known antifungals tavabarole and eubiol, and multiple steroid compounds. We also identified two structurally related compounds NSC 697923 and BAY 11-7082 (**Figure 2B**), which are phenyl vinyl sulfones. A limitation, however, is that they had some host toxicity, as measured by a significant reduction in cell mass relative to the DMSO vehicle control in Hek293 and HepG2 cells using the CellTiter-Glo assay with IC_50_ values of 252 nM and 1260 nM, respectively (**Figure 2C**). Despite this, we chose to focus on these phenyl vinyl sulfone primary hit compounds due to the specificity of the inhibition and the potential for examining structure-activity relationships and identifying new antifungal targets.

**Figure 2:**
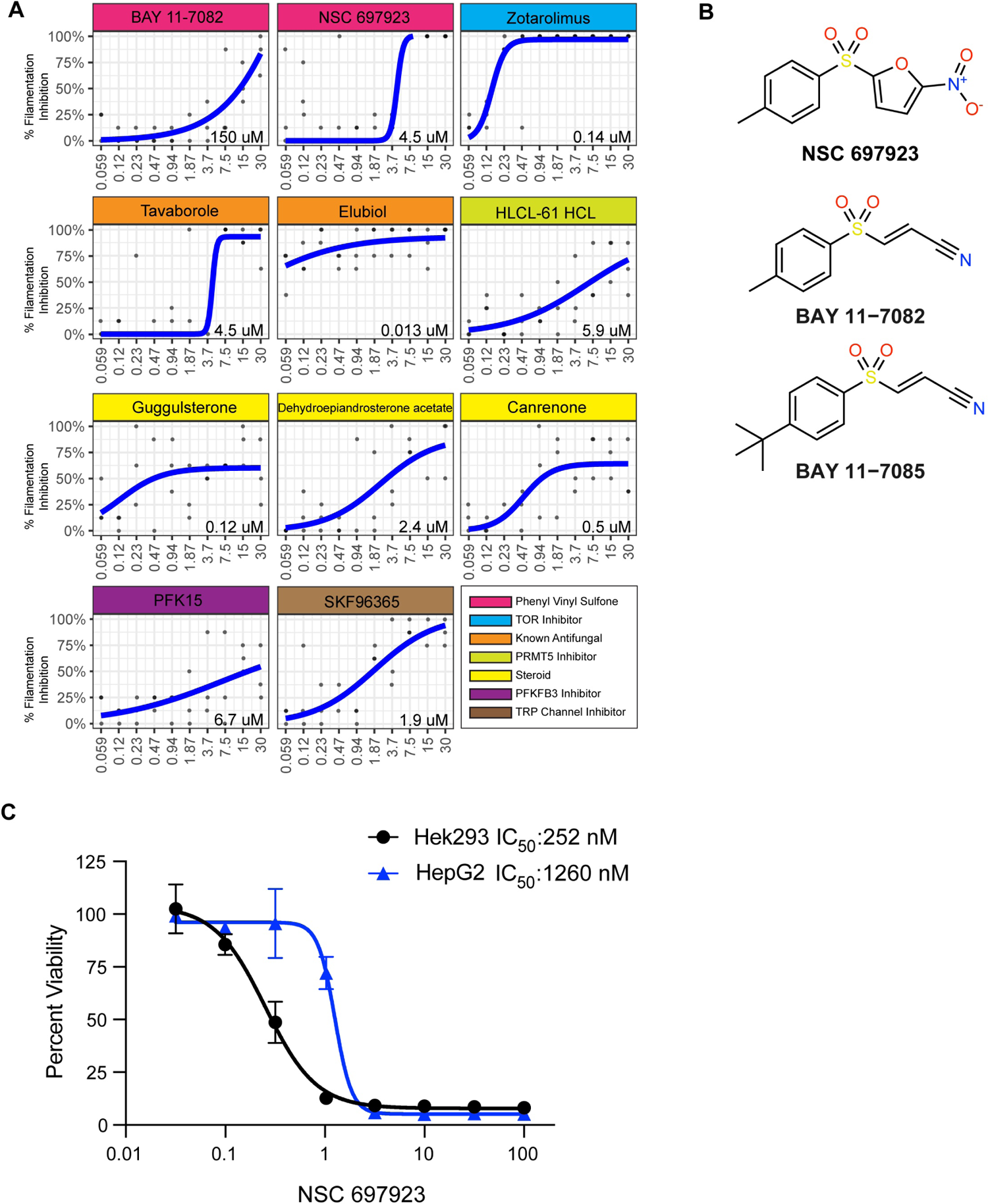
Phenyl vinyl sulfone family of compounds identified as dose-responsive filamentation inhibitors. A) Multiple screening hits displayed dose-responsive filamentation inhibition. 10-point dose-response curves were performed on potential hits from the high throughput screen and the percent filamentation inhibition was plotted. Blue line represents the best fit curve, and IC50 values were fit with 4 parameter sigmoid models using the drc R package. B) Chemical structures of phenyl vinyl sulfone family of compounds. Graphic created using ChemDraw. C) Mammalian cell toxicity of NSC 697923. Hek293 and HepG2 cells were incubated for 72 hours with an 8-point dilution series of either NSC 697923 or DMSO control and viability was measured via CellTiter-Glo luminescence. Raw luminescence was normalized to the mean counts of the vehicle control (100% viability) and empty wells (0% viability). IC50 values were determined GraphPad Prism 9.5.1 using a four-parameter inhibitor vs. response model.

### Mechanism of Action Elucidation

Our next goal was to identify the mechanism of action. Recently, the structurally related BAY 11-7082 and BAY 11-7085 compounds were shown to have efficacy against *C. albicans* and mixed culture *S. aureus* and *C. albicans* biofilms (21, 22), but the mechanism of action was still undefined. Here, we focused on NSC 697923 as it was the most potent inhibitor. NSC 697923 was initially identified in a screen for inhibition of an NF-κB signaling-responsive luciferase reporter in HEK-293T cells. The compound blocks the formation of a thioester bond between ubiquitin and the active site cysteine of the Ubc13-Uev1A E2 ligase, thus preventing NF-kB signaling. However, the ortholog of this protein is not present in *C. albicans,* suggesting that there may be another mechanism in fungi to allow for this anti-filamentation effect.

To identify a potential mechanism of action for these compounds in *C. albicans*, we employed a filamentation selection system to identify mutants that are resistant to NSC 697923. In this system, the nourseothricin resistance gene (NAT) is under control of the hyphal-specific promoter *Hwp1*. By selecting on a combination of NAT and 20 µM NSC 697923 in the presence of serum as an inducing cue, we can identify mutant strains that are specifically resistant to filamentation inhibition (Figure 3A). This approach has been successfully employed to identify genetic circuitry controlling filamentation (24).

**Figure 3:**
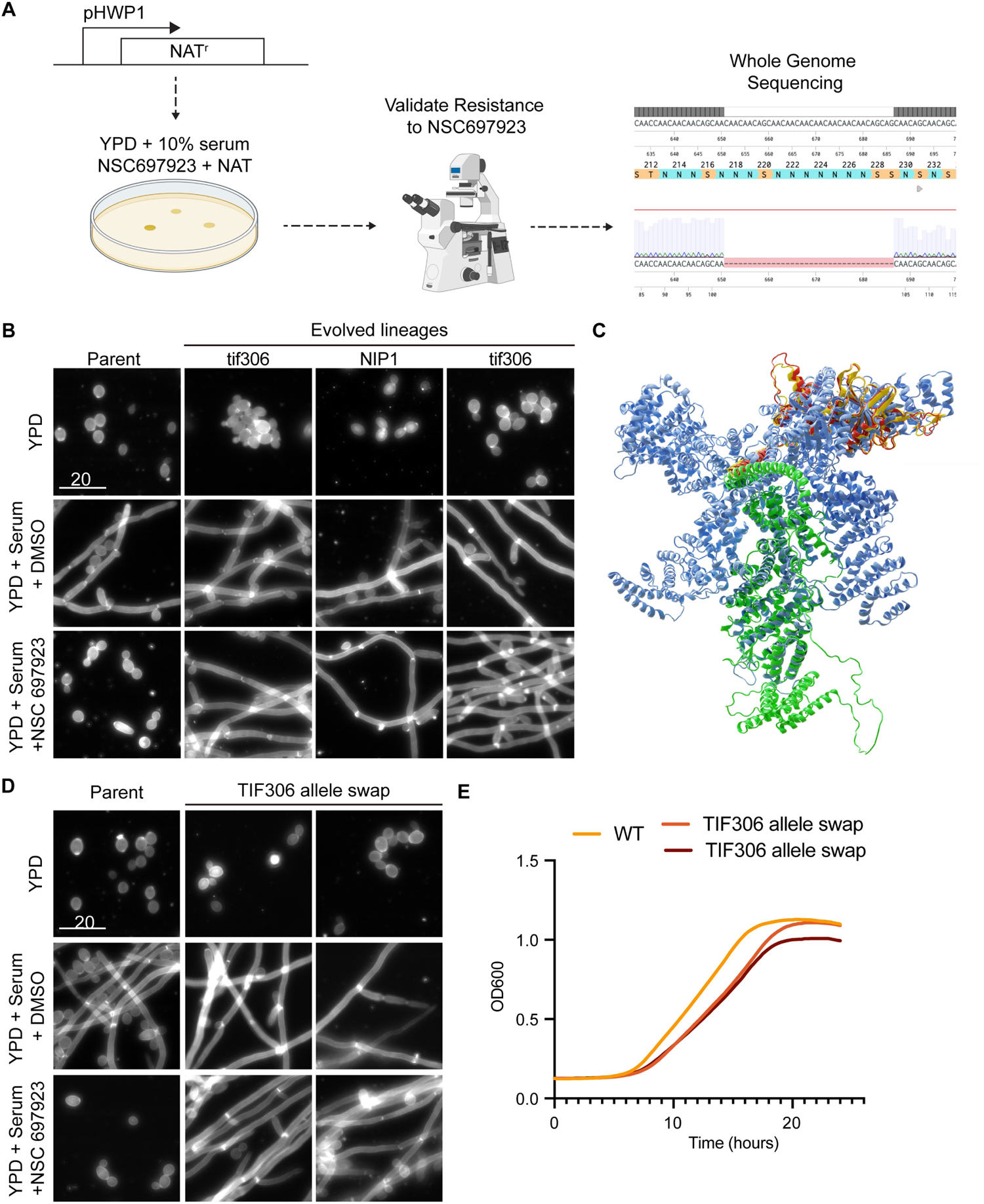
eIF3 complex identified as target of NSC 697923 and novel regulator of *C.* albicans filamentation. A) Schematic of selection regime. Selection for NSC 697923-resistant strains was driven by using a filamentation-driven NAT resistance cassette. Evolved strains were imaged to validate resistance and whole-genome sequenced for SNPs. B) 3 evolved lineages demonstrate resistance to anti-filamentation effects of NSC 697923. Images of parental and evolved lineages of *C. albicans* after incubation in YPD as a non-inducing control, YPD + 10% serum with DMSO as a solvent control, or YPD + 10% serum with 3.125 µM NSC 697923 for 24 hours. Cells were stained with calcofluor white and imaged at 40X magnification. Scale bar is 20 microns. C) Predicted protein structures of yeast eIF3 translation complex, with the *C. albicans* mutant structures in NIP1 in green, eIF4F in yellow, eIF4F mutant in red, and the mammalian reference eIF4 in light blue. Structure predicted using ColabFold v1.3.0 and visualized using PyMOL and the MoleculeNodes Blender addon. D) Mutation in TIF306 confers resistance to the filamentation inhibition of NSC 697923. Images of parental and allele-swapped strains of *C. albicans* after incubation in YPD as a non-inducing control, YPD + 10% serum with DMSO as a solvent control, or YPD + 10% serum with 3.125 µM NSC 697923 for 24 hours. Cells were stained with calcofluor white and imaged at 40X magnification. Scale bar is 20 microns. E) TIF306 allele-swapped mutant strains do not show a growth defect compared to SC5314 *C. albicans*. Cells grown 24hr in YPD for a kinetic growth assay quantified by OD600.

Using this selection regime, we identified multiple resistant colonies that were able to filament under the combination of NAT and NSC 697923. One potential mechanism of this resistance would be for the strains to be constitutively filamentous. Indeed, we identified some strains that formed hyphae even under YPD conditions, suggesting a non-specific selection for hyphal formation. However, we also identified 3 strains that showed specific resistance to NSC 697923 filamentation inhibition (Figure 3B).

To identify the genetic changes underlying this resistance phenotype, we performed whole genome sequencing of the evolved strains. We identified multiple instances of a deletion or mutation in proteins in the *eIF3* translation initiation complex; two of the strains had insertions in the *eIF3f* protein (C5_02660C or *tif306*), and one strain had an insertion in the *NIP1* protein (*eIF3c*). The *eIF3* complex is essential for translation throughout eukaryotes, but the composition of the complex varies between species (25). Additionally, *eIF3f* is a cysteine-type deubiquitinase (26); this class of enzyme is the canonical target of the vinyl sulfone drugs, suggesting that this may be the target of NSC 697923.

To evaluate a potential mechanism of the drug and resistance mutations, we aligned structures of *C. albicans* eIF3f, the mutant version of eIF3f, and the *C. albicans* NIP1 proteins, predicted using ColabFold (27), a web-based interface to AlphaFold2(28), with an orthologous experimentally determined structure of the eIF3 43S ribosomal preinitiation complex from the European Rabbit (Oryctolagus cuniculus) solved to a resolution of 6 Å through single-particle CryoEM (PDB: 5A5T) (29) (Figure 3C). *Ca* eIF3F and NIP1 have only modest sequence identity with the corresponding rabbit eIF3F and eIF3C subunits (26% and 32% sequence identity), but by leveraging the deep conservation of the core translational machinery, we found high-quality multiple-sequence alignments for each. This enabled AlphaFold2 to predict high-confidence structures (pLDDT scores of 76.9 and 70.7). Upon aligning the predicted fungal structures with the characterized mammalian complex, we found them to have close structure alignment with RMSD values of 3.3 Å over 199 of the 270 atoms, and 1.6 Å over 386 of 750 atoms, respectively, aligned using PyMOL(30). In the complex, we observe that amino acids 778-804 of eIF4F and 307-335 of NIP1 directly interact. This suggests that beyond participating in the overall function of the translation initiation, modulating the function of eIF3F or NIP1 may selectively modulate the function of the other. Interestingly, the resistant allele mutation in eFI3F modifies the length of what is predicted to be an unstructured loop. While it may be unstructured in vivo, it is also possible that this loop modulates a protein-protein interaction that is not captured in the structure of the subunit or complex.

To test whether the evolved mutations in the *eIF3* initiation complex were sufficient to confer resistance to NSC 697923, we performed allele swap experiments where we replaced the wild type allele with the putative resistance allele. Using this approach, we determined that addition of a single copy of a mutated *tif306* allele to the wild-type strain was sufficient to confer resistance to the anti-filamentation effects of NSC 697923 (Figure 3D). This provided evidence that NSC 697923 inhibits filamentation in *C. albicans* through the *eIF3* complex.

To test whether NSC 697923 has a direct effect on translation in *C. albicans*, we used a direct fluorescence-based translation assay. During active translation L-homopropargylglycine (HPG), a methionine analog containing an alkyne moiety, is incorporated into newly translated proteins and can be detected by fluorescence after copper-catalyzed click reactions with a red fluorescent Azide Plus dye. Treatment of wild type *C. albicans* with 25 µM NSC 697923 for 10 minutes resulted in an increased fluorescent signal compared to control-treated cells, demonstrating modified translation (Figure 4A, B). Notably, we observed variable translation in the control cells and uniformly high fluorescence in the NSC 697923-treated cells. In contrast, treatment of the TIF306 allele-swapped strain did not result in an increase in translation (Figure 4C, D), and there was a statistically significant although minor decrease in overall fluoresence. This provides further evidence that NSC 697923 is inhibiting hyphal formation by acting through the *eIF3* complex.

**Figure 4:**
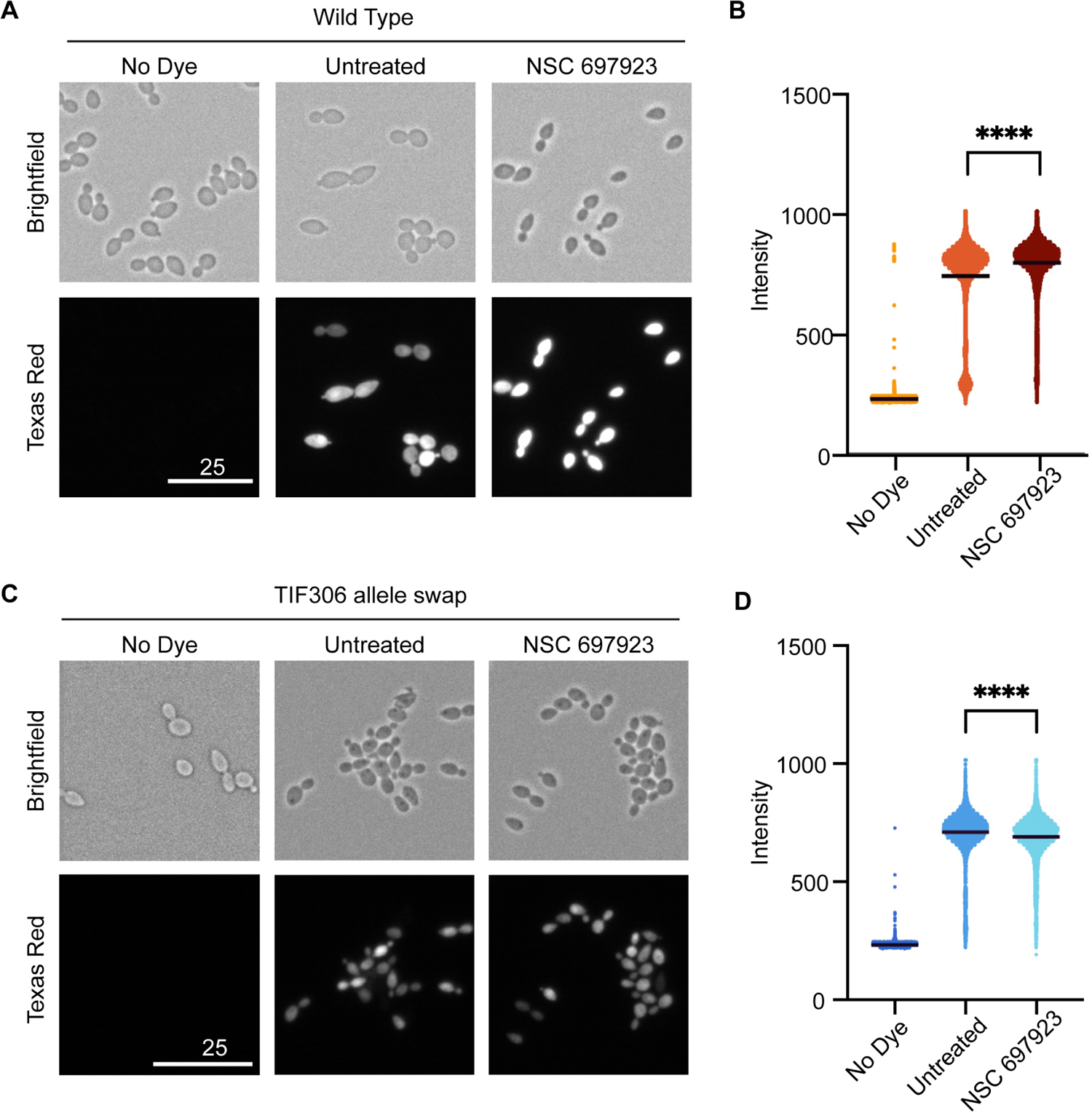
NSC 697923-treated *C. albicans* cells demonstrate modified translation. A) Use of fluorescent click-chemistry translation assay kit shows modified translation in NSC 697923-treated wild-type SC5314 *C. albicans* cells. Cells were subcultured and then treated for 10 min with 25 uM of NSC 697923. To label newly translated proteins, the l-homopropargylglycine (HPG) alkyne methionine analog was added, the cells were fixed, and the azide fluorophore was added. Cells were imaged on the Texas red channel at 40X magnification with the same exposure times to detect if translation had occurred, and with brightfield to identify cells. Scale bar indicates 25 microns. B) Fluorescence intensity results of click-chemistry translation assay of WT *C. albicans* quantified by flow cytometry on a BD Fortessa flow cytometer and analyzed using FloJo. **** indicates p-value < 0.001, student’s t-test comparing untreated and NSC 697923 treated cells. C) The TIF306 mutation precents NSC 697923 activity. Translation assays were performed as in a), using the TIF306 mutant strain. D) Flow cytometric analysis of fluorescence was performed on the cells from c), as described in b).

### Interactions with other antifungals

Combination therapies are an important strategy for increasing the efficacy of the current antifungal repertoire (5). Recently, a *C. auris-*specific translation inhibitor was discovered to enhance the efficacy of fluconazole (31). Therefore, we wanted to investigate whether NSC 697923 may act in synergy with known antifungals in *C. albicans.* To test this, we performed checkerboard assays with fluconazole, amphotericin B, and caspofungin in combination with NSC 697923. Surprisingly, we identified slight antagonism between these antifungals and NSC 697923 in *C. albicans* (Figure 5). Potentially, the increase in translation when treating with NSC 697923 results in increased resistance to known antifungals.

**Figure 5:**
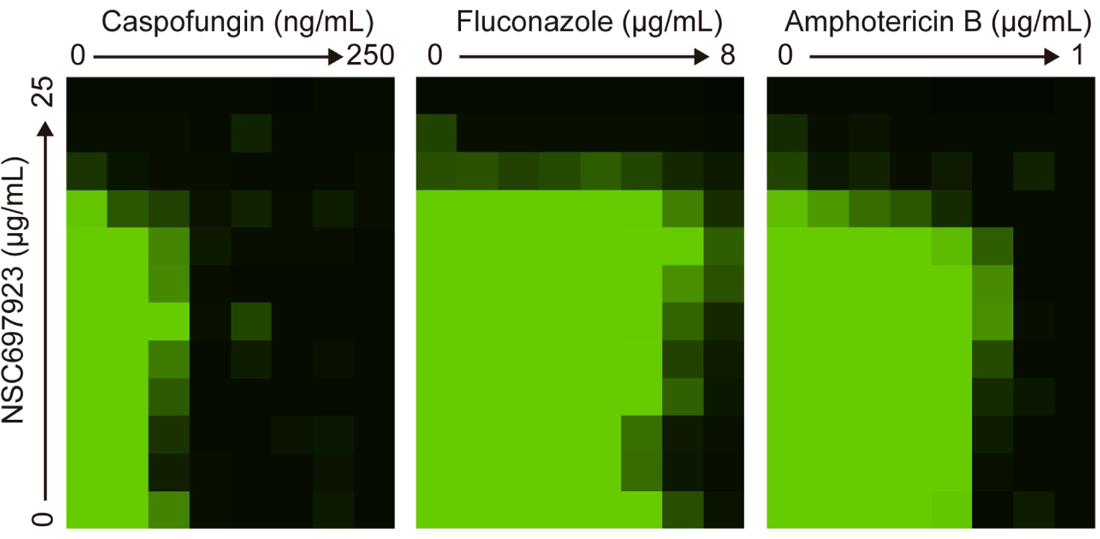
NSC 697923 shows antagonistic activity with representatives of classical antifungal therapies. A dose-response matrix was used to show the interaction between two compounds on growth in YPD at 30° C The antifungal compounds were 2-fold diluted across the X-axis, and NSC was 2-fold diluted on the Y-axis. Cells were grown for 24 hours and growth was measured by determination of the optical density at 600 nm.

## Discussion

*Candida albicans* is a fungal pathogen which is capable of causing life threatening disease, especially in immunocompromised individuals. Rising rates of antifungal resistance and limited diversity in current treatments contribute to the need for novel therapeutics to combat fungal infections, including candidemia. Antifungal resistance can evolve when selective pressure is applied to essential processes (32, 33), therefore, we aimed to target virulence factors in *C. albicans* instead of essential processes. Specifically for *C. albicans*, we targeted the morphological transition from yeast to hyphae, a known factor contributing to *Candida albicans’* ability to cause disease (8).

To find new small molecule inhibitors of filamentation in *C. albicans,* we developed and used a high-throughput microscopy screening assay to screen an FDA repurposing library of 2,017 compounds. This new assay, which built upon recent advances in morphological profiling and automated imaging analysis through CellPose and CellProfiler software, was designed to allow for screening of phenotypes besides just cell death, as many small compound libraries have already been screened for growth inhibition. The pipeline described is robust and efficient, allowing for a large number of compounds to be screened in a short amount of time, and could be applied to a explore other phenotypes, both inside and out of the field of fungi. Furthermore, this imaging-based approach is able to differentiate between fungal and mammalian cells, and for future projects it could be used to monitor the effects of small molecules while in co-culture.

In this study, we showed that one of the chemical families that was a hit in the screen, phenyl vinyl sulfones, inhibit hyphal formation in *Candida albicans* by interfering with initiation of the translation of proteins. *C. albicans* is naturally resistant to the translation inhibitor cycloheximide due to steric hindrance at the E-site (34), limiting our ability to interrogate the role of translation in this species. However, it is known that hyphal formation in *C. albicans* requires proper translation of fungal proteins (35–37). We found that the most potent phenyl-vinyl sulfone compound (NSC 697923) prevented filamentation by modifying translation through the *eIF3* complex. Furthermore, *eIF3f* also has activity as a deubiquinase (26), which is the function of the canonical target of the phenyl-vinyl sulfones in this screen. The initiation complex *eIF3* is also present in mammalian cells, which could result in off target effects if NSC 697923 was being used to treat *C. albicans* infections *in vivo.* However, targeting the translation complex may be fruitful for antifungal development, as the essentiality of members of the eiF3 complex components varies between *C. albicans* and humans (25).

While this compound was effective in targeting a virulence factor in *Candida albicans* in vitro, it likely would not be useful in a clinical setting for a few reasons. First, NSC 697923 is toxic to mammalian cells, possibly through translation modulation through the *eIF3* complex (Figure 2C). Second, at higher concentrations (>25uM), NSC 697923 was able to inhibit growth of fungi and was therefore nonspecific in targeting only hyphal formation.

Overall, we developed high-throughput screening tools that make it possible to screen additional diverse small compound libraries for new inhibitors of hyphal formation in *C. albicans*. Furthermore, the use of the pHWP1-NAT allowed us to force a non-essential function (filamentation) into temporary essentiality, which was useful in elucidating the mechanism of action in our compound after it was identified by initial screening. Future work may include medical chemistry approaches to create analogs of NSC 697923 with higher specificity. Lastly, we showed in this study that NSC 697923 demonstrated antagonistic behavior when used in tandem with the three major classes of antifungal compounds, despite NSC 697923 apparently working through a completely different method of action. It is unknown why these compounds interact in this way, leaving avenues for further exploration.

## Methods

### Strains, Reagents, and Culture Conditions

Overnight fungal cultures were grown in YPD (1% Yeast Extract, 2% Bacto Peptone, 2% Glucose) at 30°C with rotation from *C. albicans* strains archived in 50% glycerol stored at −80°C. Subcultures were grown in minimal media (2% glucose, 0.67% yeast nitrogen broth with ammonium sulfate and without amino acids) and grown at 30°C with shaking. Bay 11-7085, Bay 11-7082, and NSC 697923 were purchased from Cayman Chemical and dissolved in dimethyl sulfoxide (DMSO) solution to indicated concentrations. All strains used are available in Supplemental Table 3. For assessment of mammalian cell viability, Hek293 and HepG2 cells were both cultured in DMEM supplemented with 10% FBS, 1X Pen-Strep, and cell mass was assessed using CellTiter-Glo (Promega, Cat# E6120).

### High Throughput Screening

Overnight culture of *Eno1* promoter tagged strain CaTO31 was diluted to an OD600 of 0.03 in YPD and YPD with 20% Bovine Serum. 25 µL of dilute culture with Bovine Serum were added to columns 1-22 containing small-molecule compounds, and 25 µL of fungus in YPD without serum were added to columns 23-24 as no-drug controls in a clear-bottom 384-well plate (Grenier µclear). Cells were incubated for 5 h at 37°C and then fixed using 4% Paraformaldehyde (PFA), stained with 1% Calcofluor White (Sigma-Aldrich, catalog #18909) in PBS, and imaged on a ThermoFisher CellInsight CX5 platform using an Olympus 20X/0.45NA LUCPlanFLN objective lens microscope with 5 fields/well.

### Imaging Analysis

#### Cell painting module

To capture both yeast and hyphal forms, individual cells segmented from the images using the OmniPose algorithm(38) from the CellPose package v0.7 Nov2021(39), with parameters channels=[0,0], diameter=20, net_avg=True, agument=True, flow_threshold=10, min_size=0, rescale=True, omni=True and default parameters otherwise and saved instance masks. Images were then processed using the open-source cell segmentation software CellProfiler 4.0 (McQuin et al., 2018).

#### Scoring Cellular Phenotypes

To identify drugs that mimic the PC control, we trained a classifier to discriminate the PC and NC treated cells. To pre-process the cellular features, we first removed zero-variance and location features resulting in 384 morphological features. Then, to correct for potential differences in staining intensity, we standardized each feature on each plate using the sckit-learn StandardScalar module. We then trained an XGBoost model (v1.6.1) with default parameters over the PC and NC treated cells from plates 4 through 8 with an 80/20% train/test split. As quality control, we computed per-plate Z-prime scores. To identify important features driving discrimination, we used SHapley Additive exPlanations (SHAP), as they can handle features may have non-trivial correlation. We use the SHAP analysis implemented XGBoost library to generate SHAP diagrams and identify hi-performer features. To visualize and cluster cells based on their phenotype, we used UMAP to non-linearly embed cell-feature vectors into 2D dimensions.

The analysis pipeline, example images, and analysis code can be found at DOI: 10.5281/zenodo.7838679

To identify drugs that induce novel phenotypes, for each well we computed distance scores to each control condition. Specifically, for each well, we computed the mean cell-level feature and reduced them to 10 dimensions using SKlearn’s principal component analysis (PCA) package. Then, for each well-vector, the euclidean length and distance to each of the mean PC and NC vectors were measured.

#### Automated Dose Response Assays for filamentation

Dose-response assays were performed as previously described(40). Briefly, compounds were dispensed using an HP D300e Digital Compound Dispenser and normalized to a final DMSO concentration of 0.1% DMSO. Dose– response assays were performed in technical triplicate with 10-point/twofold dilutions. Each well was analyzed for filamentation as described above.

#### Mammalian Cell Toxicity Assay

Hek293 and HepG2 cells were plated in Corning 3675 white clear bottom 384 well plates at 3000 cells per well in 25uL and were allowed to attach overnight. Compounds were dispensed as 10mM DMSO stock solutions using a HPD300e digital dispenser and all wells were normalized to 0.2% DMSO with a top concentration of 100uM, 3.16-fold, 8-point dilution series in duplicate. After 72-hours of incubation, 25uL of the CellTiter-Glo reagent was added to each well and the luminescence signal was recorded on a BMG ClarioStar plate reader. Raw luminescence was normalized to the mean counts of the vehicle control (100% viability) and empty wells (0% viability). The replicate wells were averaged and non-linear curve fitting was performed to determine IC_50_ values in Graphpad Prism 9.5.1 using a four-parameter inhibitor vs. response model.

#### MIC assays

Standard drug susceptibility testing was evaluated by broth microdilution MIC testing in 96-well, flat-bottom microtiter plates, as previously described (41). Test compounds were dissolved in DMSO. Assays were set up in a total volume of 0.2 ml/well with twofold serial dilutions of compounds, as indicated. Growth was quantified by measuring the optical density at 600 nm (OD600) using a spectrophotometer (BioTek) at 30°C every 15 minutes with shaking for 24 hours. All strains were assessed in biological triplicate experiments with technical duplicates. Growth curves were plotted in R.

#### Selecting resistant strains

Approximately 4*10^7^ CaTO11 cells were plated on YPD containing 10% bovine serum, 20 µM of NSC 697923, and 500 µg/mL NAT. Plates were incubated at 30°C until growth of resistant colonies was observed (approx. 48hrs). Large colonies were selected and streaked to single colonies on YPD agar. To test for constitutive filamentation, each mutant strain was grown in YPD media at 30°C without a filamentation inducing cue before imaging using brightfield microscopy (20X magnification). To test for resistance to the test compound, the selected strains were grown again in YPD with 10% serum and 20 µM of each corresponding drug and observed for filamentation using brightfield microscopy (20X magnification).

#### Whole Genome Sequencing and Analysis

Genomic DNA was extracted from saturated overnight cultures, and each strain was Illumina sequenced at the Microbial Genome Sequencing Center (MiGS) as previously described (42). Variants were identified using MuTect (version 1.1.4) (43) compared with the parental CaTO11 strain, as previously described (24). Mutations identified were validated by Sanger sequencing of each region flanking the mutation (Azenta Life Sciences, Chelmsford, MA, USA). All primers are included in Supplemental Table 4.

#### eIF3 structure characterization methods

To predict the structures of eIF3F (C5_02660C_A), the NTNN mutant of Ca eIF3F, and Ca NIP1 (C4_01490W_A), we used ColabFold version 1.3.0 with building multiple sequence alignments using MMseqs2 (UniRef+Environmental) (27), using the AlphaFold2-ptm model, with 12 recycles and reporting 5 top models and default parameters otherwise. We used PyMOL to align the structure with the top pLDDT score to the mammalian eIF3 complex (PDB: 5A5T), and then rendered the overlayed structure using Blender and the MoleculeNodes addon (https://zenodo.org/badge/latestdoi/485261976).

#### Cloning/Plasmid Construction

Plasmid pTO199 was constructed using the pUC19 cloning vector, a NATflp cassette, and the *tif306* gene from CaTO197, and all pieces were amplified using primers oTO278 and oTO279, oTO765 and oTO735, and oTO763 and oTO764, respectively. The plasmid was assembled using the Codex DNA Gibson Assembly Ultra kit (Codex DNA, San Diego, CA, USA) and sequence verified by Sanger sequencing (Azenta Life Sciences, Chelmsford, MA, USA) using primer oTO531.

Plasmid pTO200 was constructed using the puc19 cloning vector, a NATflp cassette, and the *tif306* gene from CaTO199, and all pieces were amplified using primers oTO278 and oTO279, oTO765 and oTO735, and oTO763 and oTO764, respectively. The plasmid was assembled using the Codex DNA Gibson Assembly Ultra kit (Codex DNA, San Diego, CA, USA) and sequence verified by Sanger sequencing (Azenta Life Sciences, Chelmsford, MA, USA) using primer oTO531. All plasmids used are available in Supplemental Table 5.

#### Fungal Strain Construction

Transformed strains were constructed using a PCR-based transient CRISPR approach as described previously (44). Transformations used nourseothricin (NAT) as a selectable marker and were plated on YPD plates containing 150 μg/ml NAT.

To make SC4314 strains containing the *tif306* mutations from evolved strains CaTO197 and CaTO199, oTO529 and oTO865 were used to amplify the mutant allele and the NAT resistance cassette from pTO199 and pTO200 by PCR and transformed into CaTO1. Integration of the NATflp cassette and mutant *tif306* gene was tested using primers oTO2 and oTO951. For further confirmation, the *tif306* genes were PCR amplified from the transformed strain using oTO529 and oTO530 and submitted for Sanger Sequencing (Azenta Life Sciences, Chelmsford, MA, USA) with the primer oTO531.

#### Fluorescent Translation Assay

To observe translation in *C. albicans* cells, we used a Click-iT Protein Synthesis Assay Kit (Thermofisher) per the manufacturer’s instructions. WT *C. albicans* was subcultured from overnight cultures in YPD into minimal media and allowed to grow to log phase (OD600 0.4 – 0.8). After this growth, non-control cultures were treated with 25 µM NSC 697923 for 10 minutes. 50 µM HPG Reagent was used. Cells were fixed with 4% Paraformaldehyde (PFA) and permeabilized with 0.5% Triton X-100 in PBS. To quantify translation, cells were imaged on a Biotek Lionheart microscope at 40X magnification using a Texas Red channel at the same exposure time for each image. The same cells were also examined by flow cytometry on a BD Fortessa flow cytometer and analyzed using FloJo (BD Biosciences).

## Acknowledgements

This work was supported by Michigan Drug Discovery (TRO, JS), NIAID K22 (TRO), and University of Michigan Undergraduate Research Opportunities Program (KM). We acknowledge support from the University of Michigan Institute for Clinical and Health Research (MICHR) (NCATS - UL1TR002240) and its Center for Drug Repurposing (JZS). We thank all members of the O’Meara lab for helpful comments.

---

**Supplemental Figure 1: Estrogen-like compounds identified as hits in preliminary screen** Representative images of *C. albicans* from the image-based screen. Cells were stained with 1% calcofluor white, and images were taken at 20X magnification.

## Supplemental Tables

**Supplemental Table 1:** Compounds screened

**Supplemental Table 2:** Dose Response analysis

**Supplemental Table 3:** Strains used in this study

**Supplemental Table 4:** Primers used in this study

**Supplemental Table 5:** Plasmids used in this study

